# Melanopsin-dependent direct photic effects are equal to clock-driven effects in shaping the nychthemeral sleep-wake cycle

**DOI:** 10.1101/2020.02.21.952077

**Authors:** Jeffrey Hubbard, Mio Kobayashi Frisk, Elisabeth Ruppert, Jessica W. Tsai, Fanny Fuchs, Ludivine Robin-Choteau, Jana Husse, Laurent Calvel, Gregor Eichele, Paul Franken, Patrice Bourgin

## Abstract

Nychthemeral sleep-wake cycles (SWc) are known to be generated by the circadian clock in suprachiasmatic nuclei (SCN), entrained to the light-dark cycle. Light also exerts direct acute effects on sleep and waking. However, under longer photic exposure such as the 24-hour day, the precise significance of sustained direct light effects (SDLE) and circuitry involved have been neither clarified nor quantified, as disentangling them from circadian influence is difficult. Recording sleep in mice lacking a circadian pacemaker and/or melanopsin-based phototransduction, we uncovered, contrary to prevailing assumptions, that circadian-driven input shapes only half of SWc, with SDLE being equally important. SDLE were primarily mediated (>80%) through melanopsin, of which half were relayed through SCN, independent of clock function. These findings were used for a model that predicted SWc under simulated jet-lag, and revealed SDLE as a crucial mechanism influencing behavior, and should be considered for circadian/sleep disorder management and societal lighting optimization.

## Introduction

The nearly ubiquitous expression of circadian rhythmicity across species suggests that synchronizing physiology and behavior to Earth’s light-dark (LD) cycle is critical for survival and imperative for optimal functioning and health. With increasing environmental light pollution and the introduction of new technologies such as light-emitting diodes (LEDs) and connected devices, human photic behavior is increasingly uncoupled from the natural LD-cycle. The resulting perturbations in sleep-wake architecture have led to the increased prevalence of circadian disorders, insomnia, daytime somnolence, mood alteration, and poorer cognitive performance (*1–5*), stressing the need for a greater and more mechanistic understanding of the photic regulation of sleep and behavior.

Light entrains the circadian pacemaker located in the suprachiasmatic nuclei (SCN), whose output signal generates an endogenous circadian sleep-wake (SW) rhythm aligned to the environmental LD-cycle (*6*). However, light also exerts direct acute effects on sleep and waking, independent of the circadian system (*7, 8*). In nocturnal animals, such as most laboratory rodents, darkness administered for short periods (e.g. 1-hour) acutely induces waking behavior, while the same length of light exposure promotes sleep (*7*). In circadian biology these light and/or dark-dependent changes were referred to as “masking”, as they can conceal circadian-driven SW behavior. In this context, non-circadian or direct photic regulation was considered to be based upon its indirect consequences on circadian function, rather than a direct imposition on the expression of SW behavior (*9, 10*).

These non-visual effects are mediated through different photoreceptive systems and neuronal pathways. Melanopsin (Opn4), a photopigment maximally sensitive to blue light (460-480 nm), is crucial for irradiance detection (*11*). Rods and cones, the retinal photoreceptors responsible for image formation, also provide a measurement of environmental brightness (*12*). However, the degree of overlap between these two systems for regulating non-visual responses remains unclear. Non-image-forming light information is signaled from the retina to the brain through a subset of retinal ganglion cells, specifically those rendered intrinsically photosensitive (ipRGCs) by the expression of melanopsin (*13–15*). Although, these cells densely innervate the SCN via the retino-hypothalamic tract, whether this structure mediates sustained direct light effects (SDLE) is unknown. Moreover, ipRGCs send information to additional brain areas(*16–18*), including those directly implicated in the control of vigilance states such as the sleep-promoting neurons of the ventrolateral preoptic area (VLPO) (*19, 20*) or the subparaventricular zone, suggesting that direct photic regulation may play a larger role in regulating sleep than previously believed.

In recent years the development of transgenic mouse models targeting these phototransduction pathways has revealed the pronounced effects of short light and dark pulses on SW behavior (*19–21*). Moreover, we previously showed that this acute photic influence can be observed at all times of day (*20*). These observations raise the question as to whether longer periods of light/dark exposure, such as the 24-hour LD-cycle, exert sustained direct non-circadian effects, i.e. continuous acute effects that could be observed over longer periods of time. One could postulate that the complete absence of light [constant darkness (DD) experiments] would allow for the extrapolation of the influence of sustained direct photic effects, however this would not consider the possibility of altered circadian effects. Thus, under standard light-dark cycles, the contribution of direct non-circadian photic regulation in shaping the sleep-wake cycle has to date never been properly quantified, due to the difficulty in disentangling it from circadian input.

In the current study, we sought to address three related issues: (1) the quantification of the respective contribution of circadian effects and sustained direct light effects in shaping 24-hour sleep-wake distribution under standard LD-cycle, (2) the role of the SCN as a conduit for this non-circadian phototransduction of melanopsin and rods and cones, and (3) whether the quantification of circadian and SDLE allows for the prediction of the nychthemeral sleep-wake distribution under unnatural light-dark cycles, such as those experienced during jet-lag. Recording sleep in mice lacking a functional central clock and/or melanopsin-based phototransduction, we uncovered that light exerts a sustained direct influence on sleep and waking, that is primarily mediated (>80%) through melanopsin-based phototransduction, half of which passes via the SCN, implying a non-circadian function for the central structure comprising the master circadian clock. Surprisingly, our findings also reveal that SDLE and circadian effects (CE) equally contribute to shaping the nychthemeral SW cycle. To further validate our findings, these data were integrated into a model that accurately predicted 24-hour sleep-wake distribution in mice exposed to a simulated ‘jet-lag’ or trans-equatorial travel paradigms.

## Results

### Nychthemeral sleep-wake cycle amplitude is decreased by half in the absence of light or when light is equally distributed across 24-hours

In a first attempt to estimate overall light influence on the 24-hour sleep-wake cycle amplitude, we recorded sleep electrocorticograms (ECoG) in animals under DD. Here, and below, we defined the nychthemeral sleep-wake cycle (SWc) amplitude as the ratio in NREM sleep (NREMS) amounts between light (day) and dark (night) periods. Similar analyses were performed using waking or REM sleep for calculating SWc-amplitude, leading to comparable observations (see Fig. S1). Under DD, SWc-amplitude was decreased by 45% with a remaining 55% generated by the circadian drive. An alternative approximation for the overall effects of light is to analyze behavior under ultradian light/dark cycles, in which light and dark pulses are evenly distributed throughout the nychthemeron, i.e. the pulse-induced changes in sleep duration are superimposed on a circadian modulation. Mice were exposed to consecutive 1-h light and 1-h dark pulses (LD1:1), an ultradian cycle, aligned to 24-hours, which we previously used to analyze DLE according to time-of-day (*20*). An additional group of animals were exposed to consecutive pulses consisting of 3.5-h light and 3.5-h dark (LD3.5:3.5), an ultradian cycle non-aligned to 24-hour, that was specifically designed to study direct light effects independent from the circadian process (*19, 22*). In both ultradian cycles, the acute direct effects of alternating light/dark pulses were nullified (direct light effects equal to zero). Therefore, we assumed the SWc-amplitude to represent the remaining circadian effects (CE), which, remarkably, were found to be within the same range as was calculated previously under LD 12:12 (LD1:1-50 ± 11%; LD3.5:3.5-48 ± 7% vs 45% ± 6% in DD, ns; Fig. S1). Thus, this decrease by half of SWc-amplitude suggests an important role for the direct influence of light. However, the full integrity of the circadian system cannot be demonstrated under these lighting paradigms as it was unclear whether the reduction of SWc-amplitude in DD resulted from a lack of SDLE and/or altered CE, a question that our step-by-step approach below addresses.

### Sustained direct light effects are primarily mediated by melanopsin-based photoreception

Melanopsin has been shown to be the primary mediator of the acute direct influence of light on sleep and waking, with rods and cones playing only a minor role [for review see: (*7*)]. To determine whether melanopsin-based direct photic influence would be sustained for longer periods, therefore contributing to the SWc, we assessed sleep ECoG in melanopsin-deficient mice (*Opn4*^*−/−*^). Under LD12:12, mice lacking melanopsin, in comparison with wild-type animals, displayed a 1.5h reduction in NREMS, which progressively accumulated across the entirety of the 12h light phase (Fig. 1A) resulting in a 37% decrease in SWc-amplitude, similar to what we previously obtained in these mice (*20*). This reduction in NREMS also continued during the dark period, but to a lesser extent. (≈0.5h). This was interpreted as a lack of melanopsin-driven SDLE, provided that circadian rhythms were not impacted in these mice by life-long loss of Opn4. We verified this latter point by confirming SWc-amplitude under DD did not differ between mice either expressing melanopsin or not (Fig. 1B). Moreover, the accumulated NREMS loss during the 12h light phase in *Opn4*^*−/−*^ mice (Fig. 1A), was similar to the subjective light period (CT0-12) under DD in both WT and *Opn4*^*−/−*^ mice (Fig. 1C). NREMS reduction in these animals was resulted not from a circadian defect, but a lack of direct photic influence, identifying that one third (37%) of SWc-amplitude was driven by Opn4-mediated SDLE.

**Figure 1:**
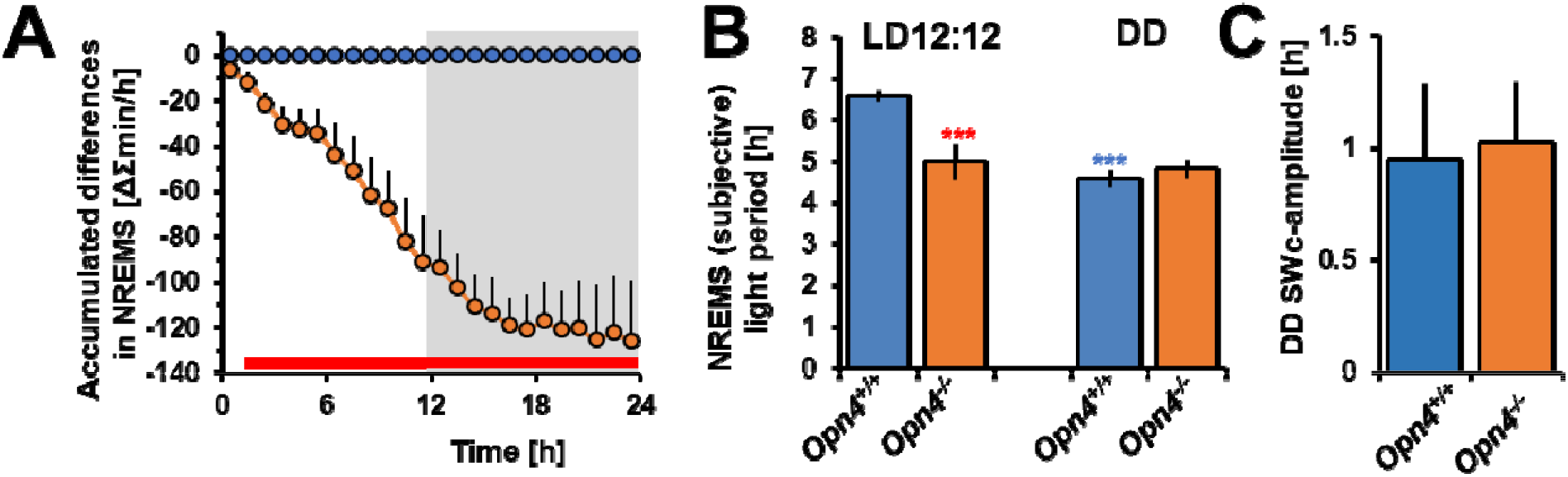
Melanopsin-based SDLE contributes to approximately one third of the SWc-amplitude. (**A**) Dynamics of the accumulated differences demonstrate that *Opn4*^*−/−*^ mice lose 2h of NREMS per day, with a maximum effect (1.5 h) during the light period (average of two consecutive days under LD12:12). (**B**) NREMS amount during light period under LD12:12 (**left**) is decreased in *Opn4*^*−/−*^ and similar to that during subjective light period under DD in both genotypes (**right**). (**C**) SWc-amplitude in *Opn4*^*+/+*^ and *Opn4*^*−/−*^ mice under DD did not differ. These observations demonstrate that sleep loss during the light period and subsequent SWc-amplitude decreases in *Opn4*^*−/−*^ mice, result from a lack of Opn4-based SDLE and not from a reduction in circadian signal. DD: Sham *Opn4*^*+/+*^ n=5; Sham *Opn4*^*−/−*^ n=7. Asterisks denote significant post-hoc differences after ANOVA between LD conditions (blue) and genotype (red).

### One quarter of the nychthemeral sleep-wake cycle is determined by SCN-independent light sustained direct effects

To calculate the proportion of the SWc-amplitude not generated by SCN signaling, we studied two groups of mice in which this structure had been disabled either through thermolytic lesioning (SCNx), or deletion of the clock gene *Bmal1* in the SCN (*Syt10*^*Cre/Cre*^*Bmal1*^*fl/−*^), which halts clock function whilst leaving the SCN physically intact (*23*). We first verified that this complete disablement of the SCN preserved the underlying phototransductive system. In SCNx mice, the lesion site was in close proximity to areas which receive direct photic input (e.g. VLPO), thus we ensured that underlying pathways were not affected through injection of an anterograde tracer, [cholera toxin subunit-B (CTb)] into the posterior chamber of the eye. We found that retinal projections were conserved in these animals to a similar extent as for sham-lesioned controls, both in cases of highly innervated brain structures such as the ventral geniculate leaflet nucleus and the superior colliculus, as well as areas less innervated but critically relevant for sleep regulation such as the VLPO (Fig. S2A-D). SCN lesioning was then verified post-mortem with AVP/DAPI whole-brain staining (Fig. S2E-H).

Our second mouse model, *Syt10*^*Cre/Cre*^*Bmal1*^*fl/−*^, has been previously shown to be arrhythmic in both SCN clock gene expression and general locomotor activity under DD, confirming an effective abolishment of the SCN clock (*23*). Additionally, to determine to what extent SCN neurons remained responsive to light, we quantified c-Fos expression, a marker of neuronal activation, in response to a 1-hour light pulse administered during the dark period [Zeitgeber time (ZT) 15-16]. This is a time-of-day when light-induced c-Fos immunoreactivity and the resulting phase-delay of the circadian rhythms of locomotor activity are known to be maximal (*20, 24*). These mice exhibited significantly diminished light-induced c-Fos expression, as compared to both control groups (*Syt10*^*Cre/Cre*^*Bmal1*^*+/−*^ and wild-type (WT) mice; Fig. 2A). A fractional conservation of light reactivity in *Syt10*^*Cre/Cre*^*Bmal1*^*fl/−*^ was observed in the retino-recipient area without transduction to the shell (more dorsal), which receives input from the core and corresponds to the clock output region (*25*). Additionally, these mice exhibited a lack of AVP immunoreactivity, a neuropeptide present in SCN cells regulating circadian behavior and interneuronal coupling (*26, 27*), whilst AVP immunoreactivity could still be detected in surrounding structures (Fig. 2Aiii,v). The absence of clock-generated signals was confirmed through sleep/wake behavioral assessment with actimetry and ECoG recordings under constant darkness (DD), which revealed a total loss of rhythmic activity as well as abolishing NREMS differences between subjective light and dark periods (Fig. 2B). Taken together, these results demonstrated the appropriateness and complementarity of two approaches models to study SDLE in the absence of CE, which controlled the putative limitations inherent to each one (i.e., lesion-induced structural damages vs. potential consequences of deleting *Bmal1* in *Syt10*-expressing cells outside the SCN).

**Figure 2:**
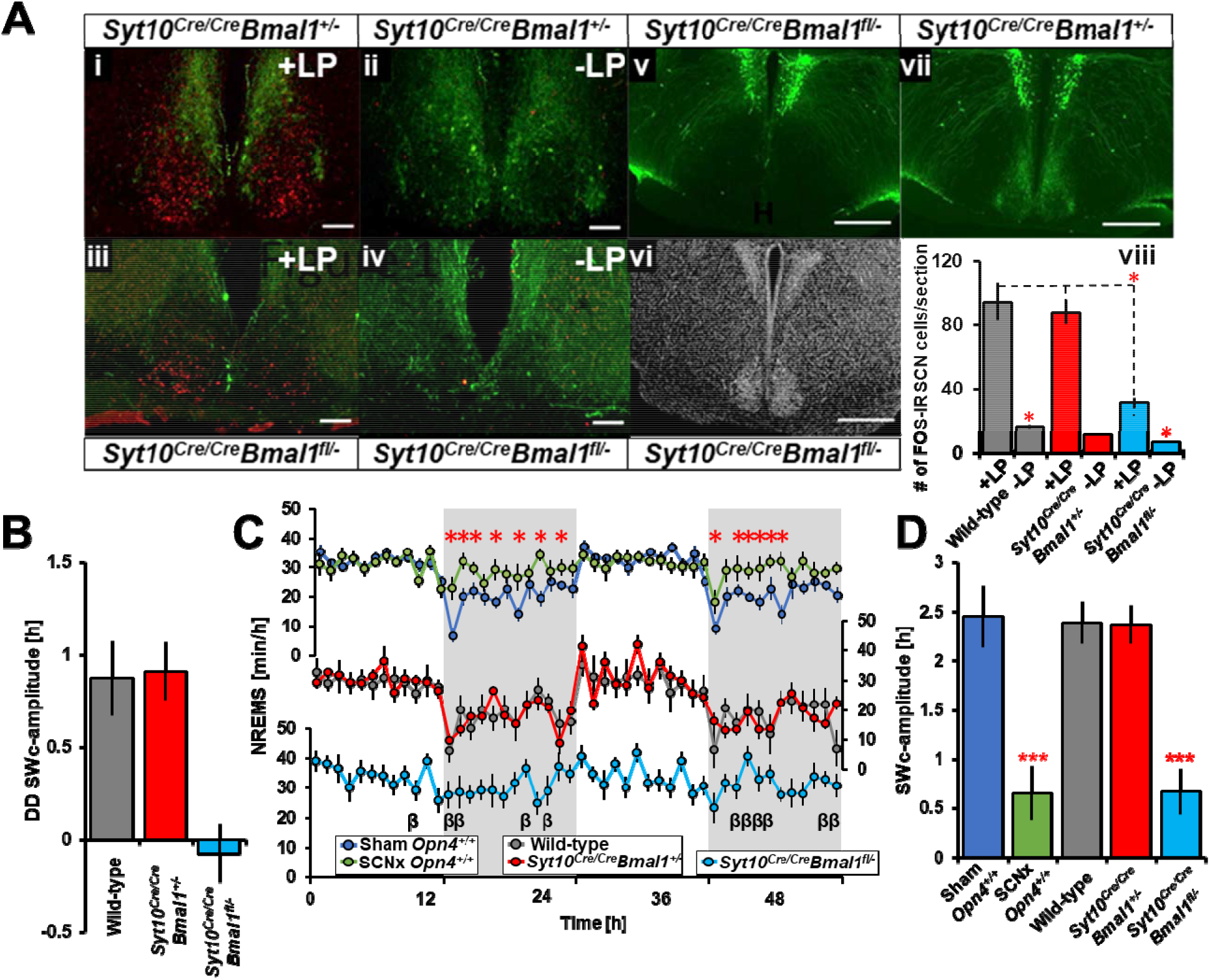
SCN-independent direct effects of light determine about one quarter of SWc amplitude. (**A)** c-FOS (red) and AVP (green) expression in the SCN. Light-pulse (+LP; at ZT15-16) induced c-FOS immunoreactivity (red) in the SCN is partially conserved in the retino-recipient (core) but absent in the clock output region (shell), in (**iii**) *Syt10*^*Cre/Cre*^*Bmal1*^*fl/−*^ in comparison to (**ii**) *Syt10*^*Cre/Cre*^*Bma1l*^*+/−*^ controls (**i**) and WT (data not shown). No c-FOS induction was seen in control conditions in the absence of a light-pulse (-LP; **ii,iv**). In *Syt10*^*Cre/Cre*^*Bmal1*^*fl/−*^ AVP (a marker of clock output, green) expression is abolished in the SCN (**v**) (that remains structurally intact; **vi**: DAPI-staining, grey), but not in surrounding areas (**iii-v**). AVP expression in the SCN is preserved in *Syt10*^*Cre/Cre*^*Bmal1*^*+/−*^ (**i,ii,vii**). Histogram represents quantification of light-induced c-FOS expression in SCN neurons (**viii**). Data are expressed as mean ± SEM. Two-way ANOVA: *P*_*light pulse*_=<0.001; *P*_*genotype*_=<0.001, post-hoc t-test: *P<0.05). No differences between controls (*Syt10*^*Cre/Cre*^*Bmal1*^*+/−*^, WT). Wild-type n=6; *Syt10*^*Cre/Cre*^*Bmal1*^*+/−*^ n=8; *Syt10*^*Cre/Cre*^*Bmal1*^*fl/−*^ n=6. Scale bars: 100µm in a-d; 500µm in e-g (**B**) Difference in NREMS amounts between subjective light and dark periods (SW-amplitude) under DD conditions in *Syt10*^*Cre/Cre*^*Bmal1*^*fl/−*^ mice and their controls. SW-amplitude of SCN-disabled mice did not significantly differ from zero. Values represent means ± SEM. (**C**) Time-course of NREMS in SCNx, *Syt10*^*Cre/Cre*^*Bmal1*^*fl/−*^ and their controls for 2 days under LD12:12 (**D**) Sleep-wake cycle amplitude (average of 2 consecutive days under LD12:12) is reduced in SCNx mice to 39 ± 13 minutes and SCN-disabled mutants (40 ± 16m), corresponding to ≈23% of that of controls. Immunohistochemistry experiments: n=3-4 per group/condition. ECoG experiments: Sham *Opn4*^*+/+*^ n=9; SCNx *Opn4*^*+/+*^ n=10; Wild-type n=7; Syt10^*Cre/Cre*^*Bmal1*^*+/−*^ n=7; Syt10^*Cre/Cre*^*Bmal1*^*fl/−*^ n=10. Psi symbols denote *Opn4* genotype post-hoc differences, delta SCN-condition, and beta *Bmal1*, respectively.

Under LD12:12 conditions, SCN-disabled mice revealed clear differences in the 24-hour time-course of NREMS (Fig. 2C). SWc amplitude under LD12:12 was approximately 2.5 hours in all control groups (Sham, wild-type, *Syt10*^*Cre/Cre*^*Bmal1*^*+/−*^). Although these models lacked a functional circadian pacemaker, we observed a remarkably similar SWc-amplitude of 23.6 and 23.2% in SCNx and *Syn10*^*Cre/Cre*^*Bmal1*^*fl/−*^, relative to controls (Fig. 2D). This indicates that light exerts a sustained direct non-circadian effect, strong enough to uphold a 24-hour organization of sleep and waking in the absence of SCN-generated circadian rhythms, and that SCN-driven signaling is responsible for three quarter of the SWc amplitude.

### The SCN acts not only as a clock, but also mediates half of the sustained direct alerting effects

Thus far we demonstrated that 77% of the SWc-amplitude depends on SCN output, and we could therefore assume this to be the result of its inherent clock function. However, we could not rule out that part of this SCN-dependent SWc-amplitude arose from a clock-independent SCN-mediation of the direct effects of light, especially given that we had previously shown the cruciality of melanopsin to this system. To separate melanopsin-based SDLE mediated through the SCN, from pathways external to these nuclei, we examined SWc-amplitude in animals lacking melanopsin photopigment (*Opn4*^*−/−*^) with or without a functional SCN (SCNx), which demonstrated that Opn4-based phototransduction accounted for 18% of the SW-amplitude through pathways outside of the SCN, and 19% within (Fig. 3A,B). Thus, our observations showed that nearly half (49%) of SDLE were mediated through SCN pathways, identifying that this structure, beyond its clock function, significantly influenced vigilance states (Fig. 3C).

**Figure 3:**
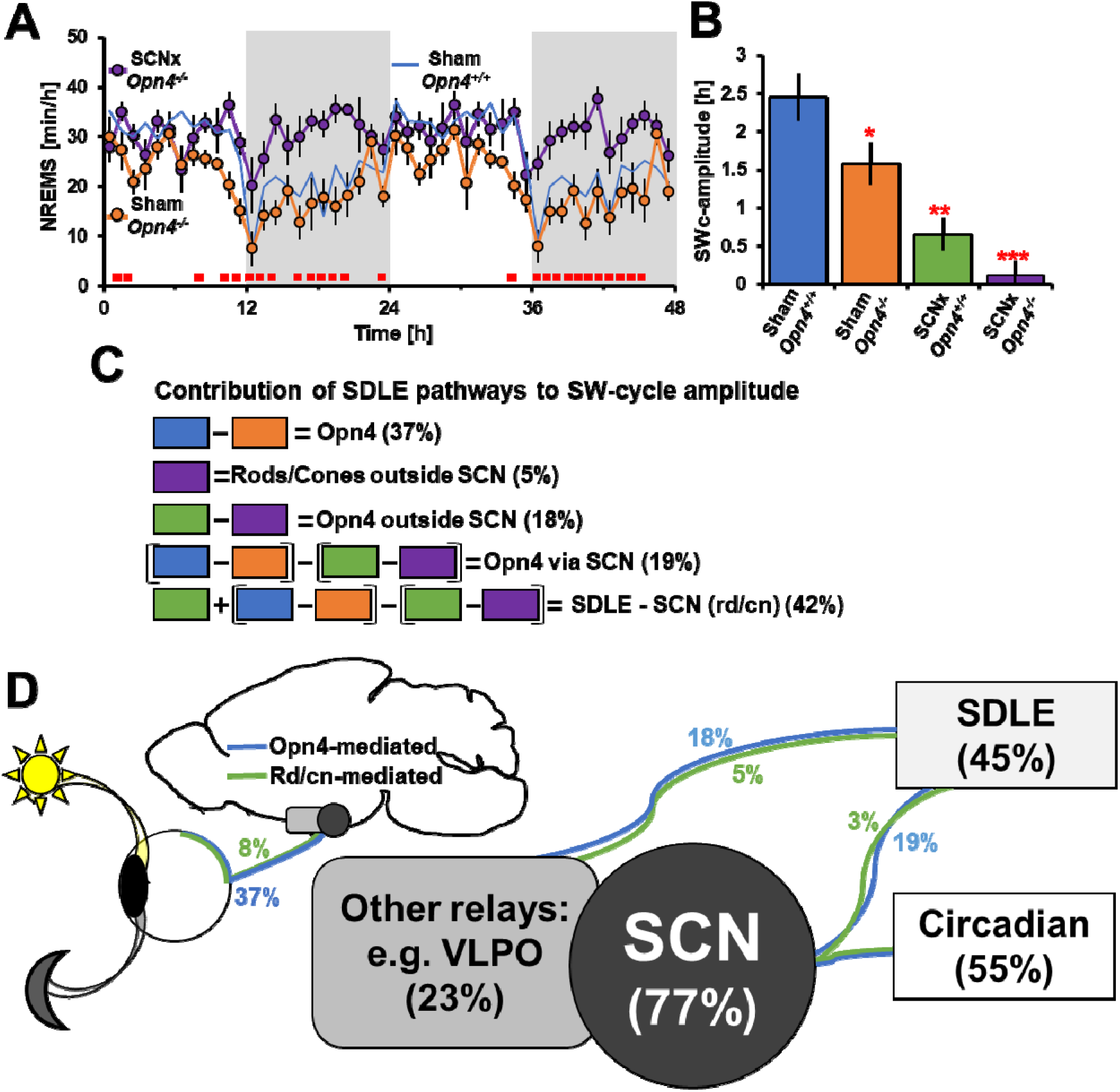
The SCN, beyond its clock function, mediates part of the sustained direct effects of light. (**A**) Time-course of NREMS during LD12:12 showed Sham and SCNx *Opn4*^*−/−*^ mice were significantly different (One-way rANOVA, group x time; *P* <0.001) as the time-course of NREMS over 24hr is flattened. (**B**) SWc-amplitude between Sham and SCNx mice with or without melanopsin. SCNx *Opn4*^*−/−*^ do not significantly differ from zero. (**C**) Intergroup differences in SW-amplitude to calculate percentage contributions to SWc-amplitude of the different pathways involved. (**D**) Respective contribution (%) of the different pathways involved in shaping the SW cycle: Contribution of CE vs SDLE (boxes), SCN vs. other brain relays and SDLE contribution mediated by melanopsin- (blue) vs. rod/cones (green) based photoreception (**see methods**). Red squares (line) in **A** represent significant post-hoc differences between groups. Asterisks represent paired t-test significance from Sham WT mice. LD12:12: Sham *Opn4*^*+/+*^ n=9; Sham *Opn4*^*−/−*^ n=7; SCNx *Opn4*^*+/+*^ n=10; SCNx *Opn4*^*−/−*^ n=8.

Interestingly, the flattened SWc-amplitude observed in SCN-ablated mice resulted mainly from a decrease in waking during the dark phase. Indeed, in nocturnal animals such as mice, darkness is known to exert a wake-promoting effect, whereas in humans, light has been shown to acutely promote alertness, with the highest efficiency of blue light centered around 460-480nm, within the spectral response peak of melanopsin (*4, 28*). Therefore, we calculated the power spectral density of the ECoG to quantify theta (7–10 Hz) and gamma (40–70 Hz) activities, known to be correlates of exploratory behavior and alertness (*29–31*). Remarkably, both were significantly decreased in each of the SCN-deficient models, specifically during the 12h-dark period of LD12:12 (Fig. S3), indicating that the alerting effect of darkness in nocturnal animals (inverted in diurnal species), is primarily signaled though the SCN.

### SDLE and CE equally contribute to shaping the 24-hr sleep-wake cycle

Although we have shown that SDLE were primarily mediated by Opn4-dependent pathways, a possible contribution of rods and cones-based photoreception could not be excluded. To address this, we calculated SWc-amplitude in SCNx *Opn4*^*−/−*^ mice, giving us an estimate of rod/cone-based SDLE mediated outside the SCN, which was found to be low (≈5%; Fig. 3B). However, these values were not significantly different from zero (one-sample signed rank test, *P*=0.22), signifying that their contribution is likely negligible. Summarizing these findings indicated that 42% of the SWc-amplitude was determined by SDLE, similar to SWc-amplitude observed in the absence of light under DD in wild-type animals (45%), and we therefore attributed this 3% difference to possible SCN-dependent rd/cn-based phototransduction (Fig. 3C).

Taken together, these results encompass different photic and circadian regulatory factors, [CE vs. SDLE, Opn4 vs. rods/cones, and SCN vs. non-SCN-mediated transmission, (summarized in Fig. 3D)], leading us to conclude that SDLE and the circadian drive are nearly equal in influencing 24-hr sleep-wake distribution. This estimate was corroborated by analysis of wild-type mice under the previously mentioned ultradian cycles, in which light and dark pulses were evenly distributed over the nychthemeral cycle, indicating approximate SWc-amplitude values of 50% and 48% for LD1:1 and LD3.5:3.5, respectively (Fig. S1). Finally, regarding other vigilance states, the contributions of CE and SDLE to REM sleep distribution were similar to those we calculated for the NREMS (Fig. S4).

### Modeling our findings predicts 24-hr sleep-wake distribution under simulated jet lag

The quantification of CE and SDLE (55 and 45%, respectively) was based solely on LD12:12 and DD conditions. We then integrated CE and SDLE contributions into a model aimed at estimating whether these respective effects could estimate overall sleep-wake distribution under other light/dark paradigms. As this was the first time SWc amplitude has been used to derive such a model, we chose two light/dark schedules simulating either transequatorial (i.e. rapid seasonal change) or transmeridian (intercontinental) travel, conditions commonly experienced in modern society, to test it. In wild-type mice exposed to a shortened photoperiod (transequatorial; LD8:16), we found the overall shape and model estimation of the nychthemeral SW-cycle, was similar to values obtained from ECoG recordings (Fig. 4A,B).

**Figure 4:**
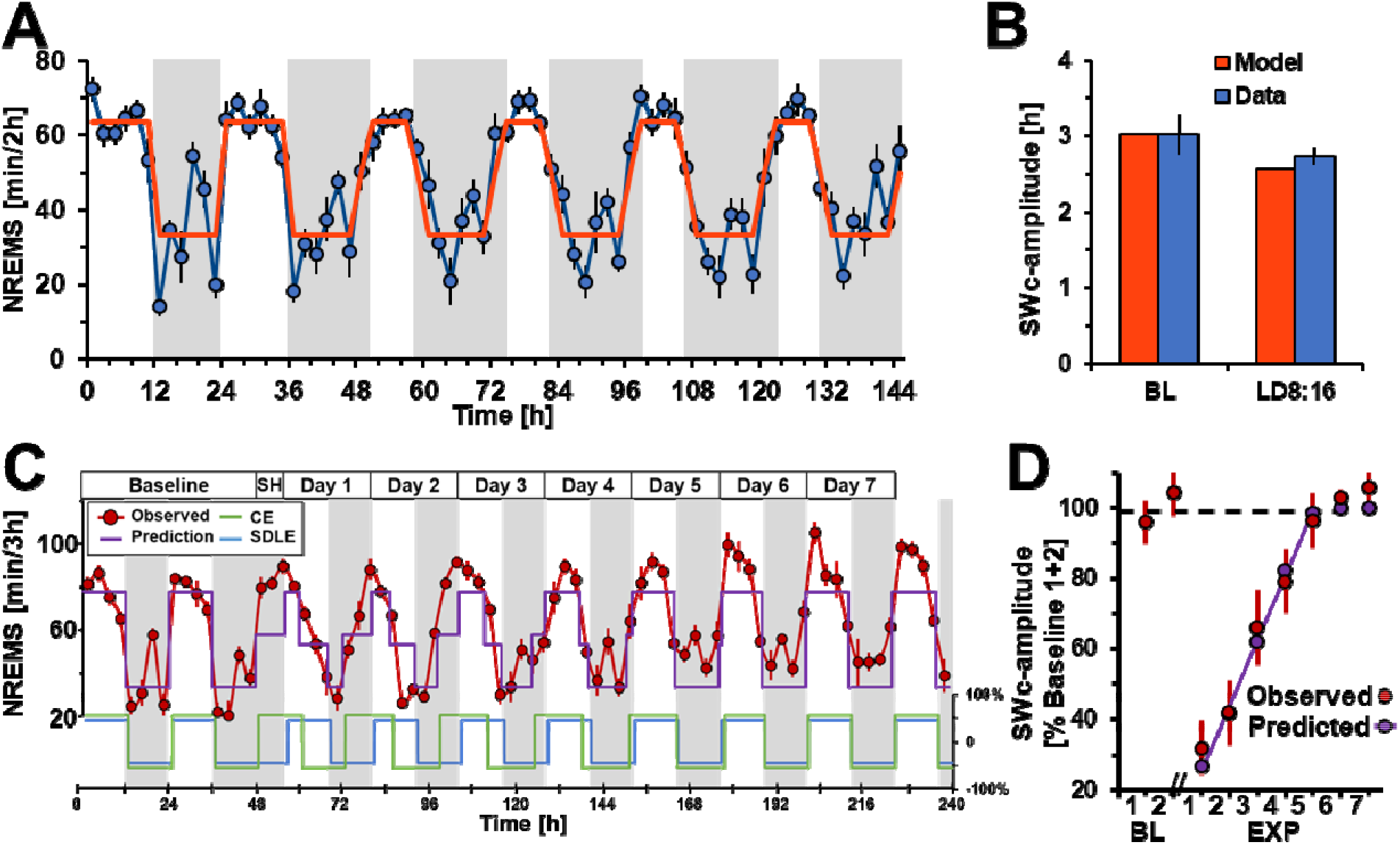
Prediction of nychthemeral NREMS distribution under simulated transequatorial and transmeridian jet-lag. (**A**) Time-course of NREMS in wild-type mice under 48-hours of LD12:12 followed by four days under 8L:16D, simulating a transequatorial travel. with observed values (blue; min/2h) and predicted values (orange; min/2h). (**B**) SWc-amplitude from prediction (orange) and from recorded animals (blue) under baseline (first 2 days under LD12:12), followed by four days under LD8:16. (**C**) Time-course of NREMS in wild-type mice under 48-hours of LD12:12 followed by seven days under a simulated 8-hour westward “jet-lag”. Predicted values of daily NREM sleep distribution (purple) based on the model integrating a shift of 1.1h/12h described in Fig. 3D, combining CE (blue) and SDLE (green). Daily NREMS distributions are obtained from sleep recordings (red) and are expressed as min/3h. SH=shift (jet-lag). (**D**) SWc-amplitude under baseline (LD12:12) and simulated 8-hour westward and “jet-lag” conditions across experimental days. A linear regression of SW-amplitude is plotted to illustrate re-synchronization to the new LD (R^2^=0.99). Values in red obtained from recorded animals perfectly fit the predicted values in purple (One-way repeated measures ANOVA, *P*_group x time_=0.28). Dashed line represents average baseline value at 100%. All values represent mean ± SEM, LD8:16: n=7; jet-lag: n=9.

Next, a separate group of mice were subjected to a simulated westward transmeridian travel with an 8-hour prolongation of the dark period, followed by a shifted LD12:12 cycle (Fig. 4C, **top**). Whereas in the first experiment we assumed the SCN output not to be affected, this condition included an additional factor, as shifting the light-dark cycle induces a phase adjustment of the clock to the new LD cycle (i.e., jetlag). Indeed, sleep recordings under this condition showed that resynchronization to the new LD12:12 cycle took approximately 96 hours (Fig. 4C). This same time-delay was observed when evaluating locomotor activity patterns, which fully resynchronized by day 5 following the jet-lag (for example, see Fig. S5). Therefore, we assumed stable entrainment to the new LD cycle across 4 days with a linear shift over this period of time of approximately 1.10h/12h, (derived from a best-fit of the observed shift, based on NREMS data). Remarkably, when integrating this constant (phase-shift of CE) into the model, the predicted 24-hour NREMS distribution (Fig. 4C, **purple line**) closely followed the observed changes in NREMS across the entirety of the experiment (11 days; Fig. 4C: **red line**), although an overall increasing trend in NREMS over the days following resynchronization could not be fully captured by the model. Furthermore, the model predicted the SWc-amplitude dynamics, substantiated by linear regression of predicted vs. observed values of these changes across subsequent days (Fig. 4D). Indeed, it captured with greatest accuracy the SWc-amplitude dynamics, which were initially attenuated as a result from CE and SDLE being out of phase and partially negating one another at certain times of the day before resynchronization (see blue and green lines in Fig. 4C, **bottom**). Taken together, these results demonstrate the robustness of this model, further validating our finding of an equal contribution of CE and SDLE in shaping nychthemeral sleep and wake distribution.

## Discussion

In this study we demonstrate, contrary to the prevailing assumption, that only half of the overall changes to SWc-amplitude are generated by a circadian signal, with SDLE being an equally important mechanism. Indeed, these findings challenge our current understanding of how photic input influences the amplitude and pattern of the SW cycle. Furthermore, we show that SDLE is capable of compensating for the absence of clock-driven influence to help maintain sleep-wake distribution aligned with external light-dark cues.

This study was designed to investigate the sustained effects of light on SW distribution, but we acknowledge that other factors potentially influence the distribution of vigilance states. In the current study, we focus only on CE and SDLE as the experiments were conducted under controlled laboratory conditions, which do not account for other environmental factors known to influence sleep and waking, Indeed, the homeostatic process, defined as an accumulation of sleep pressure with time-spent-awake and its release during sleep, is another key regulator (*32*). However, our results on SWc-amplitude and derived model, were based on 12-hour values of sleep/wake states, which likely contributes to its accuracy, with the kinetics of the homeostatic process being much faster than 12-hr periods in mice (*33*), contrary to humans. Finally, the present quantification of these CE and SDLE contributions to shaping the SWc, and the resulting modeling, is a unique resource providing a framework intended to evolve with the integration of additional environmental factors (e.g. ambient temperature, noise, etc.), which may allow to design more precise model and prediction of sleep-wake distribution at smaller time scales, also facilitating its translation to humans and other species.

Moreover, the present findings clearly redefine the role of the SCN, which should be seen not only as the orchestrator of circadian rhythms, but a critical component of the direct photic regulation of alertness. Deciphering the molecular mechanisms behind circadian and direct photic processes within SCN cells will be crucial for our understanding of how environmental signals interplay with endogenous clock-driven mechanisms for optimal physiological and behavioral functioning. Furthermore, considering that numerous advancements in chronobiology have arisen from the interpretation that observed changes in animals with SCN lesion were solely attributed to the removal of the endogenous pacemaker, the present observations should encourage revisiting some of these conclusions (*34*). In addition, our findings emphasize the role of melanopsin-based phototransduction, as compared to rods and cones, in mediating the direct effect of light on sleep-wake behavior. This corroborates current literature on the photic regulation of behavior and identifies melanopsin phototransduction-signaling as a potentially new pharmacological target for the manipulation of sleep and alertness (*7*).

This straightforward model emphasizes that optimal functioning requires CE and SDLE to be in phase with one another. Furthermore, it provides an explanation for the physiological and behavioral disturbances induced by exposure to inappropriate lighting commonly observed in today’s society, currently ascribed solely to a misalignment of the behavior relative to circadian time (*1, 2*). This could be used as a basis to optimize societal lighting and minimize light pollution, as well as its application in certain medical conditions. Indeed, current therapeutics for circadian rhythm sleep disorders (CRSD) are based upon enhancing circadian re-entrainment of the clock (*5, 35*). Our present findings, however, suggest that taking into account the direct non-circadian effects of light as a key factor of CRSD management, the insufficient consideration for SDLE possibly participating to the high rate of unsuccessful treatment (*5*). Finally, these observations could be an invaluable resource for improving other sleep disturbances, especially in facilitating adaptation to travel across time zones or to extreme latitudes, shiftwork, continuous screen exposure using modern media, or other concomitants of life in modern society.

## Material and Methods

### Animals

All experiments were performed on young adult male mice using 1) *Opn4*^−/−^ and wild-type littermate controls (*24*) on a C57Bl/6 x 129Sv genetic background and 2) *Syt10*^*Cre/Cre*^*Bmal1*^*fl/−*^ on a C57Bl/6 background (*23*), animals rendered arrhythmic through a conditional deletion of the clock gene *Bmal1* in the SCN using a Syt10 Cre-driver (*23*) (from the Max Planck Institute for Biophysical Chemistry in Göttingen, Germany). Briefly, Syt10^Cre^ mice were crossed to a mouse line carrying a conditional allele of Bmal1, allowing Cre-mediated deletion of the exon encoding the BMAL1 basic helix-loop-helix domain (*Bmal1*^*fl/fl*^). To generate experimental animals (*Syt10*^*Cre/Cre*^*Bmal1*^*fl/−*^), *Syt10*^*Cre/Cre*^*Bmal1*^*fl/+*^ females were crossed to *Syt10*^*Cre/Cre*^*Bmal1*^*fl/−*^ males. As controls, littermates of the genotype *Syt10*^*Cre/Cre*^*Bmal1*^*+/−*^ were used. To control for both the Cre-driver and the heterozygosity of *Bmal1*, we verified that *Syt10*^*Cre/Cre*^*Bmal1*^*+/−*^ and WT mice displayed similar locomotor activity, ECoG and sleep-wake patterns. All procedures complied with the local and international rules for the Care and Use of Laboratory Animals and approved by the ethical committees for animal research (Protocol N°B6748225 registered at French Research Ministry). Animals were maintained under environmentally stable conditions (LD12:12; 25 ± 0.5° C, food and water ad libitum). Genotype was validated by PCR: *Opn4*^*+/+*^ and *Opn4*^*−/−*^: primers: Mel4: 5’–TCA TCA ACC TCG CAG TCA GC–3’; Mel2: 5’–CAA AGA CAG CCC CGC AGA AG–3’; TodoNeo1: 5’–CCG CTT TTC TGG ATT CAT CGA C–3’from SIGMA life science); *Syt10*^*Cre/Cre*^*Bmal1*^*fl/−*^ and *Syt10*^*Cre/Cre*^*Bmal1*^*+/−*^ and WT: *Bmal* flox forward: 5’-ACT GGA AGT AAC TTT ATC AAA CTG-3’; *Bmal* reverse: 5’-CTG ACC AAC TTG CTA ACA ATT A-3’; *Bmal* KO forward: 5’-CTC CTA ACT TGG TTT TTG TCT GT-3’; *Syt10* forward: 5’-AGA CCT GGC AGC AGC GTC CGT TGG-3’; *Syt10* reverse: 5’-AAG ATA AGC TCC AGC CAG GAA GTC-3’; *Syt10* KI forward: 5’-GGC GAG GCA GGC CAG ATC TCC TGT G-3’. Experimenters were blind to the experimental group and condition when analyzing the data such as sleep scoring or quantification of c-Fos immunoreactivity.

## Surgery

### Lesion of the SCN

Electrolytic lesions of the SCN (SCNx) were performed under deep anesthesia (see below) and under stereotactic conditions, in a subset of mice (*Opn4*^*−/−*^, n=8) and their littermate controls (*Opn4*^*+/+*^, n=10), before electrode implantation. In order to create minimal lesions that spared surrounding brain structures, we performed a state-of-the-art 4-position electrolytic lesion of the SCN. The mice were placed in a stereotaxic instrument (Kopf Instrument), and a standard electric probe was lowered into four points of the SCN region (from zero ear bar, nose at +5°: lateral: +/−0.2 mm; antero-posterior: +3.4 and +3.6 mm; dorso-ventral: +0.95 mm). A lesion generator (Radionics Lesion Generator System) was then used to control both the temperature and voltage of the probe, for 30 seconds at each lesion site. Lesions were performed by heating the (250µm) tip of a Radionics (Burlington, MA) TCZ electrode to 55°C for 20 sec. by passing RF current from a RFG-4 lesion generator (Radionics). The same surgical and stereotaxic procedure without injected current was used for Sham control animals (the electrode was lowered into the brain using the same stereotaxic coordinates). All animals then underwent identical ECoG/EMG electrode implantation to the others (described below).

### ECoG implantation

Before undergoing implantation, mice were anesthetized with an intraperitoneal injection of ketamine 80mg/kg and xylazine 7mg/kg. Male *Opn4*^*−/−*^ (n=16), *Opn4*^*+/+*^ (n=19), *Syt10*^*Cre/Cre*^*Bmal1*^*fl/−*^ (n=10), *Syt10*^*Cre/Cre*^*Bmal1*^*+/−*^ (n=7), and *wild-type* (n=7) mice were implanted with a classical set of electrodes (two ECoG, one reference, and two EMG) in order to record vigilance states, using previously published methods(*20*). Mice were given a minimum of 14 days to recover from surgery and habituate to the sleep recording. The animals were recorded simultaneously, using commercially available hardware and software (Micromed France, SystemPLUS Evolution version 1092; Compumedics PSG4, Compumedics, Australia).

## Experimental conditions

After surgery, different groups of mice were exposed to the following light/dark conditions: (**1**) LD12:12; (**2**) constant darkness (DD); (**3**) LD1:1; (**4**) LD3.5:3.5; (**5**) Simulated transequatorial jet-lag (LD8:16); (**6**) simulated transmeridian jet-lag (8h prolongation of darkness followed by LD12:12). For anatomy experiments mice were exposed to a either a one-hour light pulse from ZT15-16, or darkness at the same time.

## Sleep and wake analysis

### Locomotor activity monitoring

General activity was monitored using a standard infrared motion detector and data were analyzed using the ClockLab software package (Actimetrics). SCNx and *Syt10*^*Cre/Cre*^*Bmal1*^*fl/−*^ mice were recorded for at least 10 days under LD12:12, followed by 2 weeks under DD to measure their free running period to confirm the loss of circadian rhythmicity. *Syt10*^*Cre/Cre*^*Bmal1*^*+/−*^ mice were also recorded under the same conditions to confirm that they display a circadian pattern of locomotor activity similar to those of WT. The actimetry recordings were also performed (in parallel to ECoG recordings).

### Scoring of sleep and ECoG power spectrum analysis

Sleep and ECoG power spectrum were analyzed and quantified according to standard criteria (*33*). ECoG and EMG signals were amplified, filtered, and analog-to-digital converted to 256Hz. The ECoG signal was then modified using a Discrete-Fourier Transform (DFT) to yield power spectra between 0 and 90 Hz (0.25Hz resolution) using a 4-s window. Any epochs containing ECoG artifacts were identified and excluded during further analyses. The behavior in each of these 4-s epochs was classified as waking, rapid-eye-movement (REM) sleep, or non-REM-sleep (NREMS) sleep by visual inspection of the ECoG and EMG signals without knowledge of the recording and genotype condition according to standard criteria. Differences between genotypes in sleep amounts were calculated by averaging time spent in each state over 5-min, 1-h, 12-h, or 24-h intervals.

For each vigilance state of the ECoG, an average spectral profile was constructed using all 4-second epochs scored with the same state. The frequency range 49-51 Hz was omitted due to power-line artifacts in some of the recordings. In NREM sleep, time-dependent changes in ECoG power for specific frequency bands, was performed for delta (0.75-4Hz). During wakefulness, theta (6-10Hz) and gamma (40-70Hz) ECoG activities were measured instead.

#### Detailed calculation of the model

The model presented in Figure 3 integrates the respective contributions of the main phototransduction pathways regulating the natural sleep-wake cycle. The contribution of each component is calculated based upon inter-group differences in amplitude of the 24-hours sleep-wake cycle (calculated as the difference in NREM sleep amounts between subjective light and dark periods). The degree of reduction in the amplitude of the sleep cycle is dependent on the phototransduction element removed in each of the groups. The specific calculations done to establish the model are shown, with the color code corresponding to each of the mouse model. The amplitude of the sleep cycle in *Opn4*^+/+^ under one day of DD immediately following LD12:12; thereby eliminating DLE i.e., allows isolating the contribution of CE (see main text). This estimate was also corroborated by analysis of the LD1:1 experiment in which the difference between averaged subjective light and subjective dark NREM sleep amounts equals the previous calculation of CE. Indeed, as mentioned in the main text, the twelve 1-hour light and 1-hour dark pulses counteract each other in the subjective light and dark period, and thus DLE = 0. After removal of CE (a), this represents DLE. The percentage passing through the SCN is represented as the difference in sleep cycle amplitude between both *Opn4*^*+/+*^ groups, with (Sham) and without an intact SCN (SCNx). The difference in sleep-wake amplitude between controls (WT and *Syt10*^*Cre/Cre*^*Bmal1*^*+/−*^ mice) and *Syt10*^*Cre/Cre*^*Bmal1*^*fl/−*^ is identical, further validating the estimation of the contribution of the SCN. The first subtraction allows calculating the Opn4 contribution to which the circadian contribution is removed to finally get the melanopsin-dependent DLE. The contribution of rods/cones-dependent DLE can be obtained from different calculations. In absence of melanopsin and SCN, the rods/cones-dependent DLE mediated outside the SCN is the only pathway that is conserved. In the absence of SCN, this intergroup difference gives us the contribution of Opn4-dependent phototransduction outside the SCN. The first subtraction allows calculating the SCN contribution, the second one the Opn4-dependent phototransduction outside the SCN, and the subtraction between both gives us Opn4-dependent DLE mediated within the SCN.

### Statistical methodology

All statistics were calculated using standard methods with Statistica (Statsoft v. 8) and graphics were generated either in SigmaPlot (Systat, v. 11) or Microsoft Excel (v. 2013). Differences in n-values between certain light/dark regimes were due to ECoG/EMG signal problems on day of recording. For ECoG spectrum analysis some animals were excluded due to the increased number of signal artifacts, which allowed for the quantification of sleep and wake distribution, but hampered Fourier signal transformation. Differences in sleep amounts and quantitative ECoG variables were determined by single-or multiple-way ANOVAs, followed by post hoc t-tests if 5% significance levels were reached. The differences in number of c-Fos positive neurons were assessed by two-way (light and genotype conditions) ANOVA, followed by post hoc t-tests.

### Retinal tract tracing

After completion of the recording protocol, SCNx and Sham animals (Sham *Opn4*^*+/+*^ n=9; Sham *Opn4*^*−/−*^ n=7; SCNx *Opn4*^*+/+*^ n=10; SCNx *Opn4*^*−/−*^ n=8; Wild-type n=7; *Syt10*^*Cre/Cre*^*Bmal1*^*+/−*^ n=6; *Syt10*^*Cre/Cre*^*Bmal1*^*fl/−*^ n=8) were anaesthetized (Isoflurane 2.5%, applied during all procedure) and 1µl of the anterograde tracer cholera toxin subunit B (CtB; 1%, dissolved in 50 mM PBS, pH 7.4) was injected into the vitreous over 2 min. After 48 hours animals were perfused, and the histology procedure was processed as described below.

## Anatomical studies

### Immunostaining

After completion of the protocol, mice were deeply anesthetized with CO2 and perfused with 0.1 M PBS pH 7.4 followed by transcardial fixation for 15 min with 4% paraformaldehyde in PBS, pH 7.4. In order to study c-Fos immunoreactivity in response to light in *Syt10*^*Cre/Cre*^*Bmal1*^*fl/−*^ (n=3), *Syt10*^*Cre/Cre*^*Bmal1*^*+/−*^ (n=4) and WT (n=3) animals the sacrifice occurred at the conclusion of a 1h light pulse administered at ZT15 and without light pulse as a negative control. SCNx and sham animals were sacrificed 48 hours after intraocular injection of CtB (see above). To control for SCN lesion size and conservation of retino-cerebral projections, immunohistochemistry was used in SCNx and Sham-lesioned animals to stain arginine vasopressin (AVP), 4’,6-diamidino-2-phenylindole (DAPI) and CtB. To evaluate SCN cells reactivity to light in *Syt10*^*Cre/Cre*^*Bmal*^*fl/−*^ and their controls, c-Fos expression was quantified in the SCN with AVP staining. The procedure was carried out as described previously(*20*). After dehydration in 30% sucrose for 48h the brains were frozen and cut in a freezing microtome. Free-floating slices, brought to room temperature, were rinsed 3 times (5 minutes each) in 0.1 M phosphate-buffered saline (PBS, pH 7.0); incubated in 5% normal donkey serum (Normal Donkey Serum 017-000-121 Jackson ImmunoResearch) and 0.25% Triton X-100 (TX) in PBS for 1 hour; and then incubated overnight at 4°C with primary antibodies (rabbit anti-c-Fos IgG, SC52 SantaCruz Biotechnology, diluted 1:1,000; guinea pig anti-AVP T-5018 Bachem, 1:2000, goat pAb anti-cholera toxin B subunit Calbiochem© 227040 1:1000) in a solution containing 0.25% TX in PBS. The following day, sections were rinsed 3 times (10 minutes each) in PBS before incubation with secondary antibodies (goat anti-rabbit IgG Alexa 555 Invitrogen 10373021 1:500; donkey anti-guinea pig conjugated Cy5 Jackson ImmunoResearch 706-175-148 1:200; donkey anti-goat conjugated Alexa 555 Invitrogen A21432 1:500) in a solution containing 0.25% TX in PBS for 2 hours. They were then rinsed 3 times (10 minutes each) in PBS and finally mounted on slides, allowed to air-dry at room temperature, and cover-slipped with mounting solution (glycerol 50%, 0.1 M PBS 50%, DAPI 1:500 Sigma-Aldrich D9542). Omission of the primary antibody abolished all staining (control).

### Photomicrographs and quantification

Immunostainings were analyzed using a microscope equipped with appropriate filter settings for detecting Cy5, Alexa 555 and DAPI. Fluorescence images were obtained via a non-confocal microscope (DMRXA2, Leica Microsystems) equipped with Metamorph v 2.1.39 (Olympus, Ballerup, Denmark). Light microscopy images were grabbed with a Leica DC200 camera using Leica DC200 software (Leica, Cambridge, UK). The software program Image J was used to fusion the images and the image editing software Microsoft publisher was used to combine the obtained images into panels. Reference of the various brain structures was made according to the Franklin and Paxinos atlas “Mouse brain in stereotaxic coordinates” (*36*).

For quantification of c-Fos-immunoreactivity to light, the number of c-Fos-positive neurons in the SCN was counted by two independent experimenters blind to experimental conditions, in an area of 300 x 500 µm at levels corresponding to bregma −0.30mm to −0.70mm (rostral to caudal part of the SCN) (Fig. 1C). Approximately 9 brain sections from each animal containing the SCN (with AVP co-staining) were analyzed and c-Fos-positive cells were visually counted using a grid on each image. C-Fos immunoreactivity in response to light was analyzed using two-way ANOVA (light pulse and genotype conditions) followed by post-hoc t-tests.

## Data and Code availability

All data and code used for the analysis presented in this paper is available from the authors upon reasonable request.

## Acknowledgements

We would like to thank D. Ciocca and S. Reibel-Fosset -Chronobiotron-UMS3415 for their support with the mice colony and their advice on experiments and planning. This work was supported by grants, from Alsace-biovalley, France Région Grand-Est - Strasbourg Eurométropole (CUS) and from Fondation Adiral. Je.H. was supported by a fellowship of the French Ministry for Research and Higher Education and Fondation Adiral.

## Author Contributions

The study was designed by Je.H., P.F., and P.B.; EEG and actimetry experiments were performed by Je.H., M.K-F., E.R., J.T., L.Ch, F.F., L.Ca., Ja.H. E.R., and L.Ch, performed the anatomy experiments. Je.H., M.F., J.T., and F.F., scored sleep/wake ECoG. Je.H. analyzed the data. Je.H., G.E., M.K-F. and P.B. wrote the manuscript. The authors declare no competing interests.

---

**Supplementary Figure 1 (S1):**
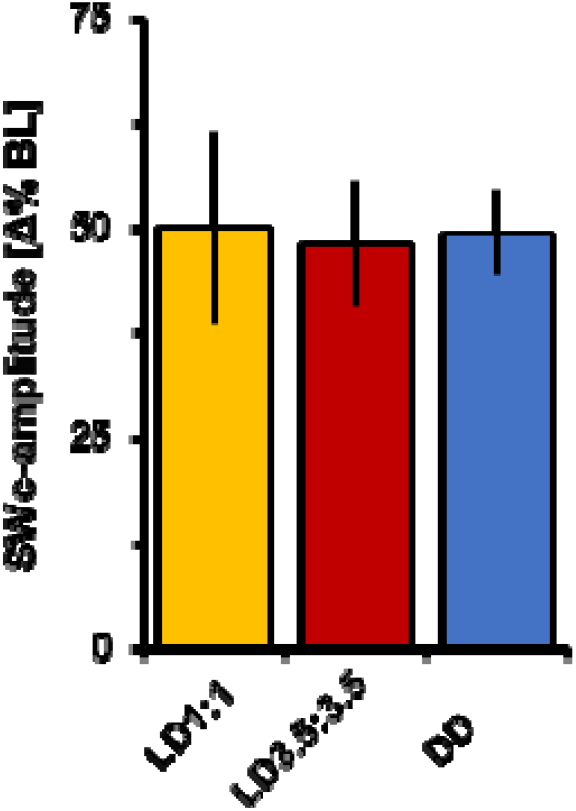
Sustained direct light effects are modified during ultradian cycles and constant darkness in wild-type mice. Wild-type mice exposed to either ultradian (LD1:1, LD3.5:3.5) or constant darkness, show reductions in SWc-amplitude of approximately 50% when compared to baseline conditions. LD1:1 n=9; LD3.5:3.5 n=6; DD n=5.

**Supplementary Figure 2 (S2):**
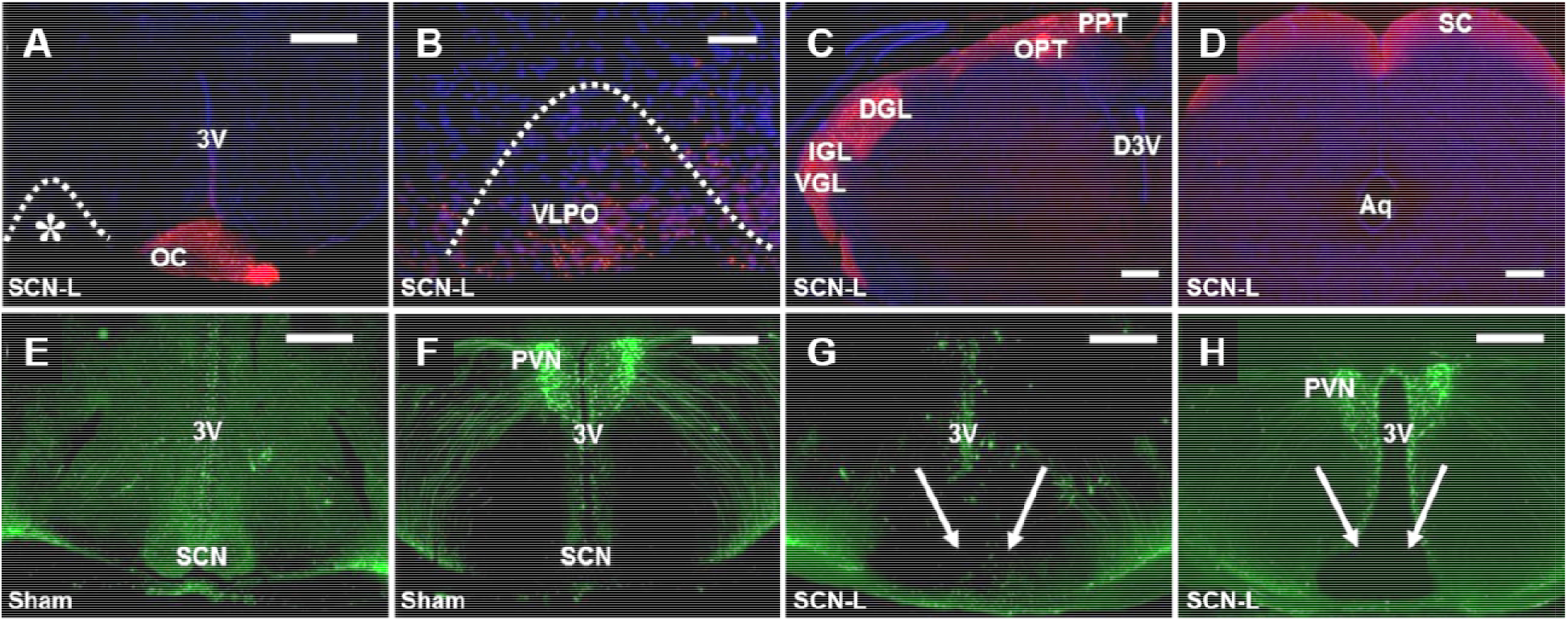
Retinal projections to the brain are preserved in SCN-lesioned animals. **Top:** Staining CTb (red) and DAPI (blue) in an SCNx (SCN-lesioned) mouse at the level of **(A)** the optic chiasm (OC); (**B**) VLPO (Enlargement of VLPO fibers from * in **a**); **(C)** geniculate leaflets (DGL, IGL, VGL), and pretectal nuclei (OPT, PPT), and **(D)** superior colliculus (SC). Retinal projections to the brain are conserved in similar proportions to those observed in sham controls. **Bottom:** Coronal sections at the (**E, G**) mid and (**F, H**) more caudal levels to the SCN in a (**E,F**) Sham and (**G,H**) SCN-L mouse stained for AVP (green). Note the complete removal of the SCN (white arrows, compared to **E**). Sham: Sham-lesioned. DGL/IGL/VGL: dorsal/inter/ventral geniculate leaflet, O/PPT: olivary /posterior pretectum, 3V: third ventricle, PVN: paraventricular nucleus. Scale bars: 500µm (**A,E-H);** 20µm (**B**) and 400µm (**D**). n=3-4 per group.

**Supplementary Figure (S3):**
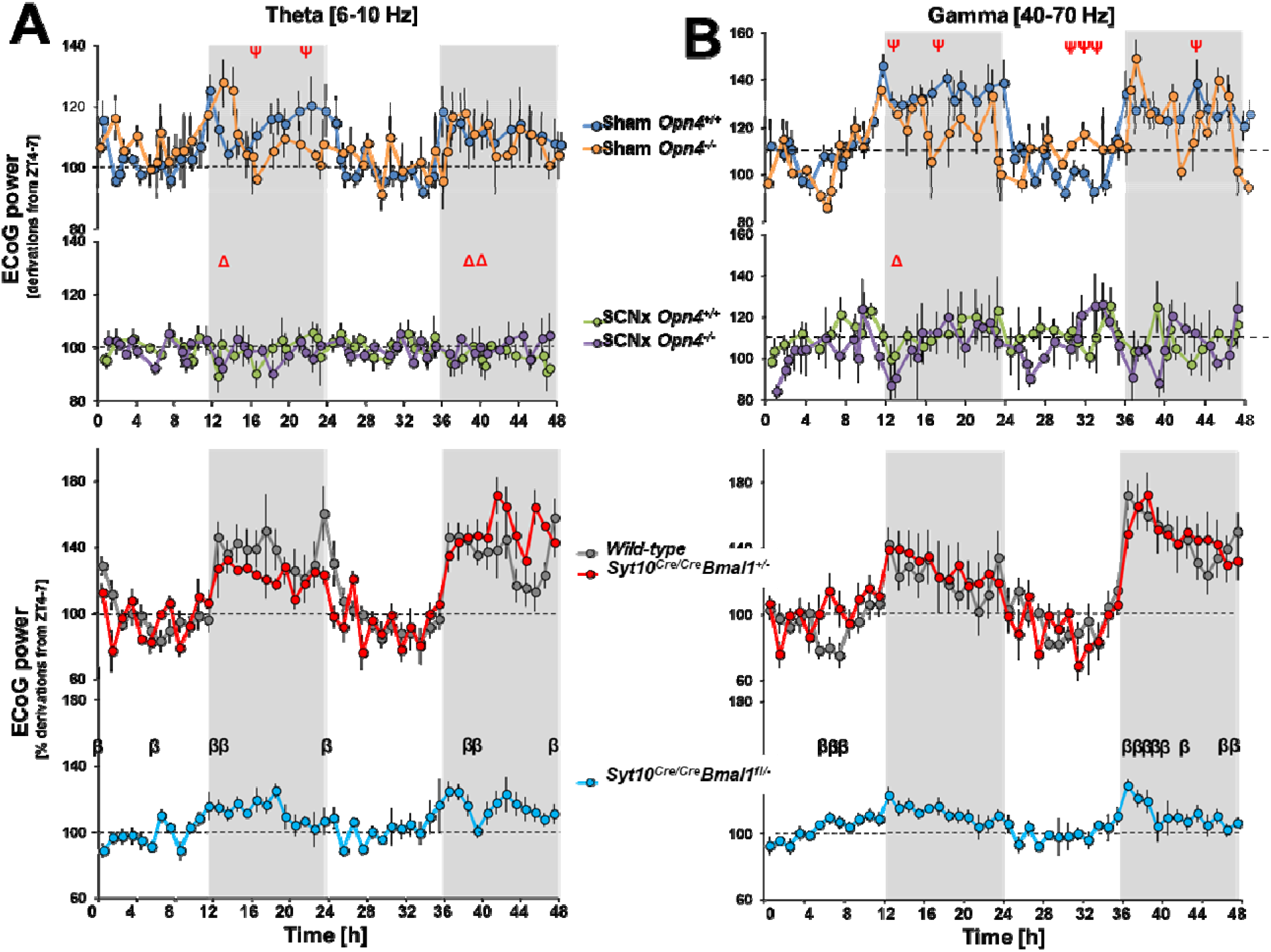
Darkness exerts a sustained alerting effect under LD12:12 which is dependent on the SCN. Darkness promotes theta and gamma ECoG activities under LD12:12 which is dependent on the SCN. Quantification of theta (**A**; 6–10 Hz) and gamma (**B**; 40–70 Hz) power spectra in waking ECoG during 2 consecutive days under LD12:12. Values (mean ± SEM) are normalized against the lowest values during the 24h period from ZT4-7. The three control groups (Sham *Opn4*^*+/+*^, wild-type, *Syt10*^*Cre/Cre*^*Bmal1*^*+/−*^, displayed higher amounts of ECoG theta/gamma activities during the 12-hour dark period (top: *P*=0.01 bottom *P*=0.003). The dark-associated promotion of ECoG theta and gamma activities was attenuated in the absence of melanopsin (*Top*: theta: *P*_genotype_<0.001; gamma: *P*_genotype_=0.011) and dramatically flattened in absence of a functional SCN [*Top*: theta: *P*_SCN-lesion_=0.002; gamma: *P*_SCN-lesion_<0.0001; *Bottom:* theta: *P*_SCN-deficient_<0.0001; gamma: *P*_SCN-deficient_<0.001; *Top:* 3-way rANOVA; genotype x SCN-lesion x time-course; *P<0.001*; *bottom:* 2-way rANOVA; SCN-deficient x time-course; *P<0.001* followed by post-hoc tests (*P*<0.05) showing significance for genotype and SCN condition depending on the time of day. No significant differences were observed during the subjective light period. ECoG power was normalized against the period of lowest theta and gamma power (ZT4-7) of each respective day (Sham *Opn4^+/+^ n=8;* Sham *Opn4^−/−^ n=6;* SCNx *Opn4^+/+^ n=5;* SCNx *Opn4^−/−^ n=5*; wild-type *n=6; Syt10^Cre/Cre^Bmal1^+/−^ n=5; Syt10^Cre/Cre^Bmal1^fl/−^ n=8*). Psi symbols denote *Opn4* genotype post-hoc differences, delta SCN-condition, and beta *Bmal1*, respectively.

**Supplementary Figure (S4):**
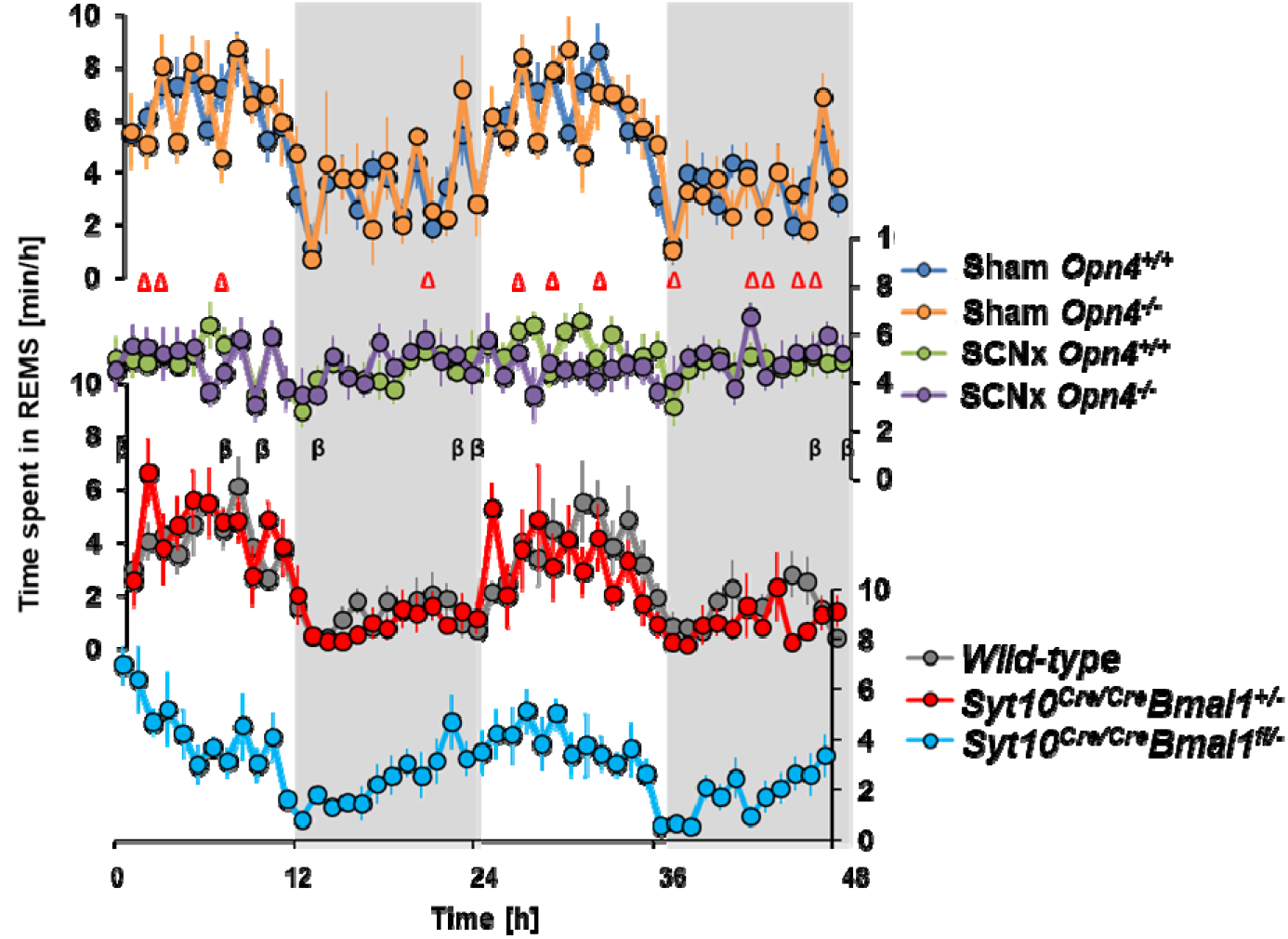
REMS cycle is flattened in absence of a functional SCN. Time-course of REMS for two consecutive days under LD12:12 is similar to that of NREMS (*P*_*time-course*_<0.0001) (Figure 1A,2D). The amplitude of the light-dark REMS cycle is flattened in the absence of a functional SCN (Top: *P*_*SCN -lesion x time-course*_<0.0001; Bottom: *P*_*SCN-deficient x time-course*_=0.003). The removal of melanopsin does not significantly affect its time-course. Control groups were not significantly different from one another. All values represent mean ± SEM. Statistics: 3-way ANOVAs followed by post-hoc *t-*tests: red delta: SCN lesion difference; black beta: genotype (SCN deficient) difference. Sham *Opn4*^*+/+*^ n=9; Sham *Opn4*^*−/−*^ n=7; SCNx *Opn4*^*+/+*^ n=10; SCNx *Opn4*^*−/−*^ n=8; Wild-type n=7; Syt10^*Cre/Cre*^*Bmal1*^*+/−*^ n=7; Syt10^*Cre/Cre*^*Bmal1*^*fl/−*^ n=10.

**Supplementary Figure (S5):**
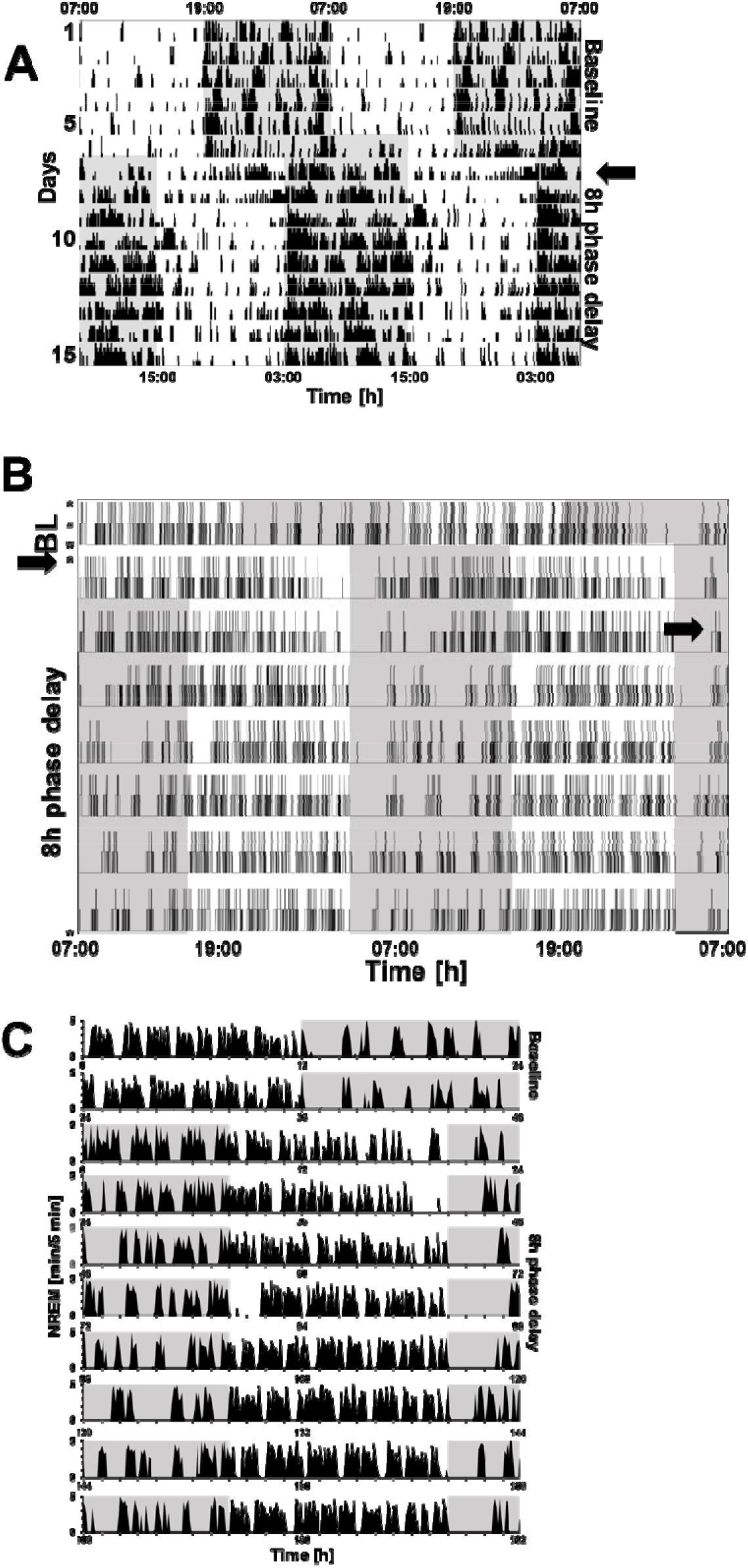
Daily expression of locomotor activity and sleep and waking in mice exposed to a simulated westward travel (8h jet-lag) Representative **(A)** double-plotted actograms of locomotor activity, **(B)** double-plotted hypnograms of sleep-wake states (W: wake, N: NREMS, R: REMS), and **(C)** single-plotted overview of wakefulness expressed per 5 minutes intervals of one animal that re-entrained under simulated “jet lag” condition. The day of the shift is indicated (→). Time is given as elapsed rather than ZT due to the phase shift of the LD cycle (Baseline: light: 07:00am-07:00pm; dark: 07:00pm-07:00am; after a phase-delay of 8-hours to simulate westward jet-lag: light: 03:00pm-03:00am; dark: 03:00am-03:00pm). Note the complete resynchronization after 4 days to the new LD condition. **(A)** Activity counts per 5-minutes are indicated in solid black. Six days of baseline are recorded followed by 9 days following jet-lag. **(B)** Hypnogram of the same mouse (ECoG/EMG recordings across the entirety of the experiment); wake (no line), NREMS (middle line), and REM sleep (upper line). (**C**) NREMS per 5 minutes in one mouse. Results are consistent with what is seen in (A). BL: Baseline.

## References

1. T. A. Bedrosian, R. J. Nelson, Influence of the modern light environment on mood. Mol Psychiatry 18, 751–757 (2013).

2. J. Falbe et al., Sleep duration, restfulness, and screens in the sleep environment. Pediatrics 135, e367–375 (2015).

3. K. M. Stephenson, C. M. Schroder, G. Bertschy, P. Bourgin, Complex interaction of circadian and non-circadian effects of light on mood: shedding new light on an old story. Sleep Med Rev 16, 445–454 (2012).

4. G. Vandewalle, P. Maquet, D. J. Dijk, Light as a modulator of cognitive brain function. Trends Cogn Sci 13, 429–438 (2009).

5. U. Kilic, J. Hubbard, P. Bourgin, in Sleep medicine textbook C. Bassetti, Z. Dogas, P. Peigneux, Eds. (European sleep research society, 2014).

6. A. A. Borbely, Effects of light on sleep and activity rhythms. Prog Neurobiol 10, 1–31 (1978).

7. J. Hubbard, E. Ruppert, C. M. Gropp, P. Bourgin, Non-circadian direct effects of light on sleep and alertness: lessons from transgenic mouse models. Sleep Med Rev 17, 445–452 (2013).

8. C. Cajochen, Alerting effects of light. Sleep Med Rev 11, 453–464 (2007).

9. N. Mrosovsky, Masking: history, definitions, and measurement. Chronobiol Int 16, 415–429 (1999).

10. W. J. Rietveld, D. S. Minors, J. M. Waterhouse, Circadian rhythms and masking: an overview. Chronobiol Int 10, 306–312 (1993).

11. R. J. Lucas, M. S. Freedman, M. Munoz, J. M. Garcia-Fernandez, R. G. Foster, Regulation of the mammalian pineal by non-rod, non-cone, ocular photoreceptors. Science 284, 505–507 (1999).

12. G. S. Lall et al., Distinct contributions of rod, cone, and melanopsin photoreceptors to encoding irradiance. Neuron 66, 417–428 (2010).

13. D. M. Berson, F. A. Dunn, M. Takao, Phototransduction by retinal ganglion cells that set the circadian clock. Science 295, 1070–1073 (2002).

14. S. Hattar, H. W. Liao, M. Takao, D. M. Berson, K. W. Yau, Melanopsin-containing retinal ganglion cells: architecture, projections, and intrinsic photosensitivity. Science 295, 1065–1070 (2002).

15. I. Provencio, M. D. Rollag, A. M. Castrucci, Photoreceptive net in the mammalian retina. This mesh of cells may explain how some blind mice can still tell day from night. Nature 415, 493 (2002).

16. J. Hannibal, J. Fahrenkrug, Target areas innervated by PACAP-immunoreactive retinal ganglion cells. Cell Tissue Res 316, 99–113 (2004).

17. J. Hannibal et al., Central projections of intrinsically photosensitive retinal ganglion cells in the macaque monkey. The Journal of comparative neurology 522, 2231–2248 (2014).

18. S. Hattar et al., Central projections of melanopsin-expressing retinal ganglion cells in the mouse. The Journal of comparative neurology 497, 326–349 (2006).

19. D. Lupi, H. Oster, S. Thompson, R. G. Foster, The acute light-induction of sleep is mediated by OPN4-based photoreception. Nature neuroscience 11, 1068–1073 (2008).

20. J. W. Tsai et al., Melanopsin as a sleep modulator: circadian gating of the direct effects of light on sleep and altered sleep homeostasis in Opn4(-/-) mice. PLoS Biol 7, e1000125 (2009).

21. C. Altimus et al., Rods-cones and melanopsin detect light and dark to modulate sleep independent of image formation. Proceedings of the National Academy of Sciences 105, 19998–20003 (2008).

22. T. A. LeGates et al., Aberrant light directly impairs mood and learning through melanopsin-expressing neurons. Nature 491, 594–598 (2012).

23. J. Husse, X. Zhou, A. Shostak, H. Oster, G. Eichele, Synaptotagmin10-Cre, a driver to disrupt clock genes in the SCN. Journal of biological rhythms 26, 379–389 (2011).

24. N. F. Ruby et al., Role of Melanopsin in Circadian Responses to Light. Science 298, 2211–2213 (2002).

25. J. A. Evans et al., Shell neurons of the master circadian clock coordinate the phase of tissue clocks throughout the brain and body. BMC Biol 13, 43 (2015).

26. X. Jin et al., A molecular mechanism regulating rhythmic output from the suprachiasmatic circadian clock. Cell 96, 57–68 (1999).

27. M. Mieda et al., Cellular clocks in AVP neurons of the SCN are critical for interneuronal coupling regulating circadian behavior rhythm. Neuron 85, 1103–1116 (2015).

28. S. A. Rahman et al., Diurnal spectral sensitivity of the acute alerting effects of light. Sleep 37, 271–281 (2014).

29. G. Buzsaki, E. I. Moser, Memory, navigation and theta rhythm in the hippocampal-entorhinal system. Nature neuroscience 16, 130–138 (2013).

30. C. S. Herrmann, M. H. Munk, A. K. Engel, Cognitive functions of gamma-band activity: memory match and utilization. Trends Cogn Sci 8, 347–355 (2004).

31. X. J. Wang, Neurophysiological and computational principles of cortical rhythms in cognition. Physiological reviews 90, 1195–1268 (2010).

32. A. A. Borbely, A two process model of sleep regulation. Hum Neurobiol 1, 195–204 (1982).

33. P. Franken, D. Chollet, M. Tafti, The homeostatic regulation of sleep need is under genetic control. The Journal of neuroscience: the official journal of the Society for Neuroscience 21, 2610–2621 (2001).

34. F. K. Stephan, I. Zucker, Circadian rhythms in drinking behavior and locomotor activity of rats are eliminated by hypothalamic lesions. Proceedings of the National Academy of Sciences 69, 1583–1586 (1972).

35. T. I. Morgenthaler et al., Practice parameters for the clinical evaluation and treatment of circadian rhythm sleep disorders. An American Academy of Sleep Medicine report. Sleep 30, 1445–1459 (2007).

36. G. Paxinos, K. B. Franklin, Paxinos and Franklin’s the mouse brain in stereotaxic coordinates. (Academic press, 2019).

